# The Landscape of Mutations in Human Fumarate Hydratase

**DOI:** 10.1101/852392

**Authors:** David Shorthouse, Michael W J Hall, Benjamin A Hall

**Affiliations:** MRC Cancer Unit, University of Cambridge, Cambridge, CB2 0XZ, UK; Wellcome Trust Sanger Institute, Hinxton, CB10 1SA, UK

**Keywords:** Mutation, Mutational Screen, Fumarate Hydratase, Molecular Dynamics, Metabolomics

## Abstract

Fumarate Hydratase (FH) is an enzyme of the citric acid (TCA) cycle that is responsible for reversibly catalysing the conversion between fumarate and malate. FH loss and subsequent buildup of the oncometabolite fumarate causes hereditary leiomyomatosis and renal cell carcinoma. We explore the mutational landscape of FH in silico, predict the functional effects of many already detected mutations, and categorise detected but un-characterised mutations in human populations. Using mutational energy predicting tools such as Rosetta and FoldX we accurately predict mutations and mutational hotspots with high disruptive capability. Furthermore, through performing molecular dynamics simulations we show that hinge regions of the protein can be stabilized or destabilized by mutations, with new mechanistic implications of the consequences on the binding affinity of the enzyme for its substrates. Finally, we categorise all potential mutations in FH into functional groups, and predict which known mutations in the human population are loss-of-function, and therefore predispose patients to papillary renal carcinoma – we validate our findings through analysis of metabolomics data of characterized cell lines.

## INTRODUCTION

Fumarate hydratase (FH) is a member of the tricarboxylic acid (TCA) cycle occurring in the mitochondria, and enzymatically metabolises fumarate within the cytosol. FH activity in the cell is responsible for the reversible conversion of the metabolite fumarate into malate, and the knockout or mutational inactivation of FH in kidneys is linked to an oncogenically-associated buildup of fumarate(1, 2). As a result the enzyme FH has been described as a tumor suppressor, and fumarate as part of a novel classification of molecules named “oncometabolites”. Precisely how the buildup of fumarate can be oncogenic is unknown, but recent work points towards suppression of DNA repair responses, EMT, and promotion of mitotic entry upon fumarate buildup(3–5).

Understanding the effects of mutations on the activity and assembly of FH is of importance for the understanding and stratification of germline mutations in FH, which can predispose patients with a single mutated or deleted allele to hereditary leiomyomatosis and renal cell cancer (HLRCC) upon mutational inactivation of their remaining wild-type copy(6, 7). Previous work has identified mutants linked with inherited and de-novo FH-related conditions, including cancer(8) – most notably, the FH mutation database represents a comprehensive list of mutations and their effects, if known, on FH activity(9).

In recent years numerous methods have been developed for estimating the effects of single point mutations (SNPs) on the stability of a protein structure in silico. Notable methods include FoldX(10, 11), which uses an empirical force field to predict the alterations in a protein induced by mutation, and methods included as part of the Rosetta suite(12, 13), which uses Monte-Carlo based dynamics to predict energetic effects of mutations. Additionally, molecular dynamics can be used to more comprehensively investigate mutant protein structure, though at significantly higher computational cost. With the advent of high-throughput methods such as CRISPR screening, and larger projects being undertaken to screen populations for mutations and disease, coupled with large-scale disease-focussed data generating projects such as The Cancer Genome Atlas (TCGA)(14) and the International Cancer Genome Consortium (ICGC)(15), the number and diversity of mutations being implicated in disease is rapidly expanding. Whilst methods to attempt to sift functionally relevant mutations from synonymous to detect highly mutated genes exist in the form of statistical tests such as DN/DS(16), mutsig(17), and oncodrive(18), including some methods that take into account structure of the protein such as Rhapsody(19), there is scope for detailed, structure-informed, chemically aware methods to classify mutations, including those not yet observed, into Loss-of-Function (LOF) and benign categories.

Here we computationally screen and classify every potential mutation in the available fumarate hydratase structure to study the landscape of potential mutations. We consider the structural and biological implications of each mutation, and thus can predict mechanistic details of every potential mutant. We confirm that our method predicts known functionally relevant mutations, and predict from existing databases of mutations which have an unknown effect, which of them will be damaging to the activity of FH. Overall we predict that 66% of all mutations to FH influence activity or assembly. We further validate our predictions through studying the Cancer Cell Line Encyclopaedia (CCLE)(20, 21) and show that previously unstudied mutations that we predict to be damaging to the function of FH result in altered metabolite levels expected from disruption to the activity of FH.

## RESULTS

### Evidence of mutational clustering in FH

Human FH is formed as a homotetramer of subunits generated from the *FH* gene. Each subunit contains 3 domains, Domain 1, Domain 2, and Domain 3 (D1, D2, and D3 respectively) (**Fig 1 A**). D1 is formed from residues in the range 49-188, D2 is formed from residues in the range 189-439, and D3 from residues in the range 440-510. The full functional protein is an assembly of 4 subunits and contains 4 identical binding pockets made of interactions between 3 subunits (**Fig 1 B**). There are two proposed regions of importance for catalysis of the fumarate/malate conversion; Site A, the known active site (hereafter referred to as the binding site), and Site B, a region of proposed but unknown functional importance(22, 23). For this study we chose to only include the known catalytic site, Site A, defined as residues HIS176, ASN182, SER186, SER187, ASN188, THR234, HIS235, LYS371, VAL372, ASN373, and GLU378 (**Fig 1 C**). We do not consider Site B due to the unknown and conflicting evidence surrounding its importance. For this study we chose to focus on the crystal structure 5upp(24), which covers residues 49-510 of the 510 residue protein assembled into a homotetramer.

**Figure 1:**
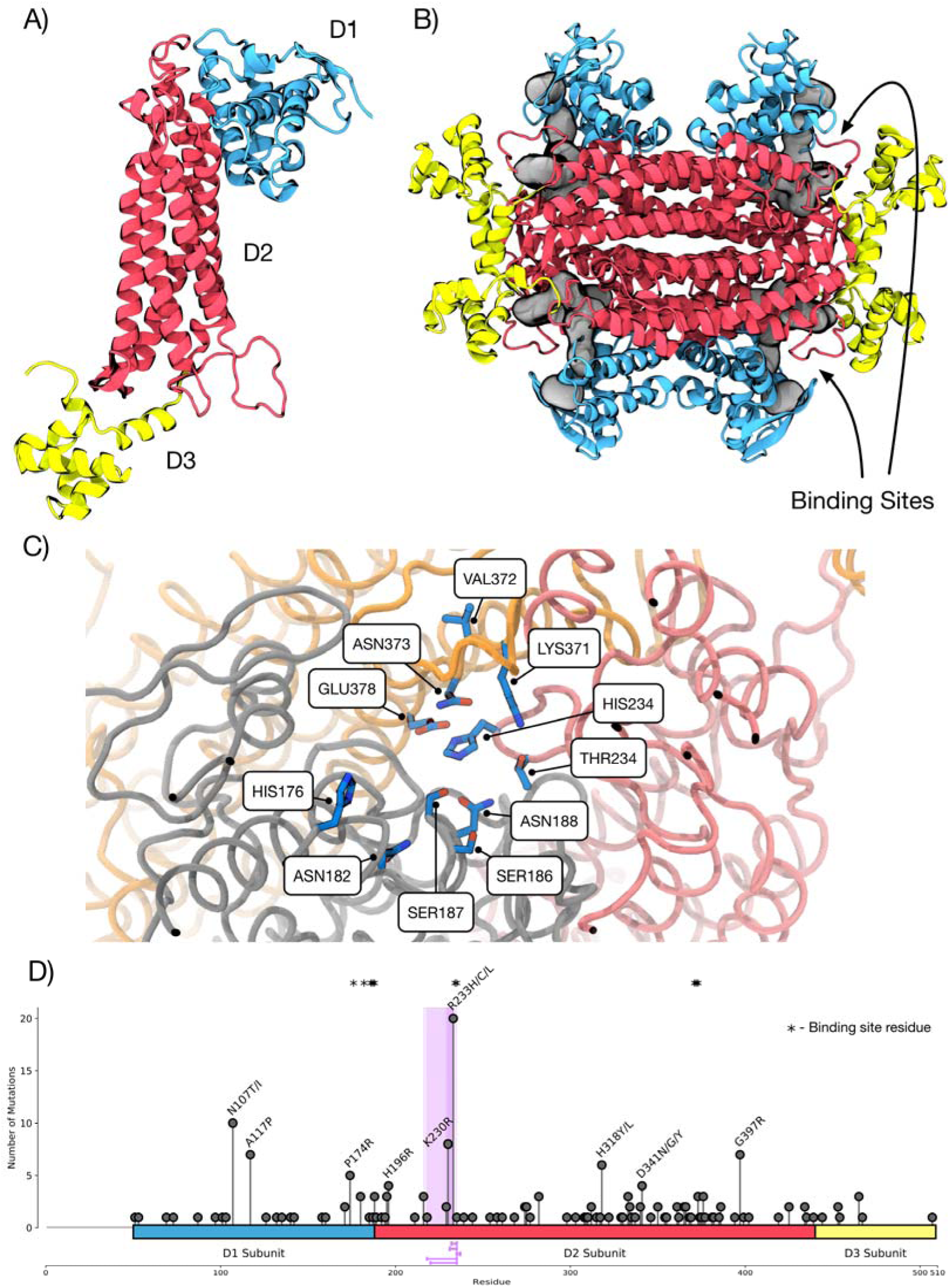
Structure and current mutations in Fumarate Hydratase. A) Structure of a single subunit of FH showing the D1, D2, and D3 regions. B) Structure of an assembled homotetramer of FH. Binding sites are highlighted and made up on an interface between 3 subunits. C) Close up of the binding site of FH showing the residues involved in catalytic activity. D) Mutational spectrum of non-benign SNPs in FH. D1, D2, and D3 regions are highlighted in blue, red, and yellow respectively. Stars indicate residues involved in catalytic activity that make up the binding site of FH. Purple highlight and lines represent the top 5 mutational clusters as calculated by the NMC algorithm.

To study mutations known or suspected to have roles in human disease, we investigated the Fumarate Hydratase Mutation Database(9), which contains 378 mutations, including 113 that are distinct missense, at the time of this study. The Fumarate Hydratase Mutation Database attempts to pool all observed mutations in FH, including those that are benign, and a large number of mutations have no clinical or functional annotation. Mutations that are not known to be benign (i.e those either labelled as loss-of-function, or those which are uncharacterised) are shown in **Fig 1 D**.

We applied the NMC clustering method to look for clustering of mutations across the 1D sequence of the protein(25). We chose to include the top 5 predicted clusters, ranked by significance, and with a size less than 50 residues long. We find the most significant clusters are all within the region of the more prevalent mutations in residues 230 and 233, indicating that this region is statistically highly over mutated, and potentially a mutationally vulnerable site.

### Classification of mutations by proximity to the binding site and protein hinges

Residues of the catalytic site in FH have been previously identified as essential for the conversion of fumarate to malate. We define binding site-associated residues as those with alpha-carbons (CA) within 6 Å of the CA of any binding site residue. Mutations this close to the binding site are likely to disrupt the assembly of the binding site, and we classify them as potentially disruptive. A significant number of known mutations are binding-site adjacent (**Fig 2 A**) and significantly mutated.

**Figure 2:**
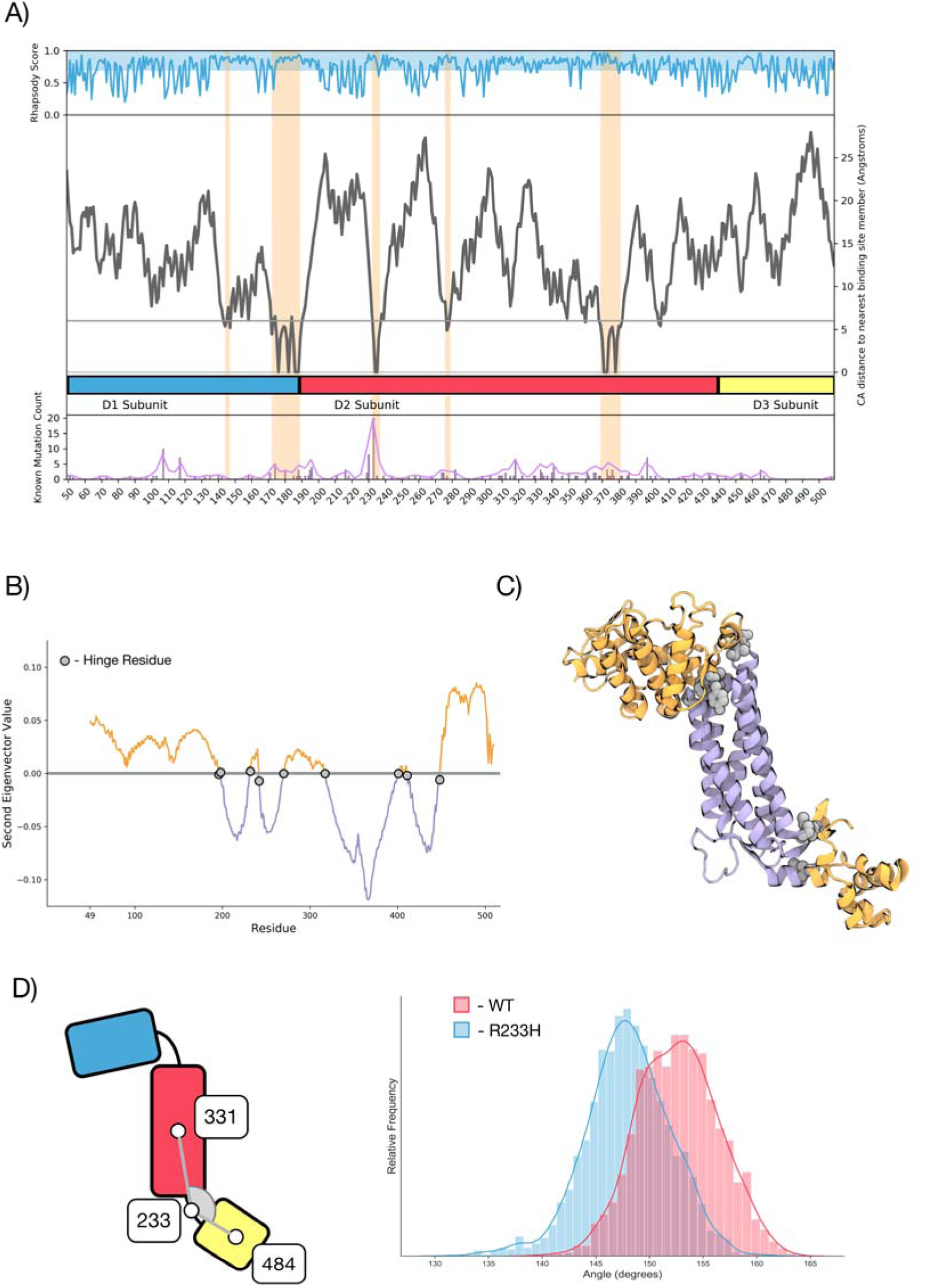
Mutations can be categorised on proximity to functional regions of FH. A) Alpha carbon (CA) distance from a binding site residue. Shown is: Top: average rhapsody score for each residue, Middle: distance of each residue from a binding site residue by CA distance, Bottom: mutational frequency for each residue. Orange highlights show some regions have high rhapsody scores, low binding site distance, and high mutational frequency. B) Second normal mode eigenvectors per residue for a single subunit of FH. Residues with an eigenvector above the line are moving generally opposed to those with an eigenvector below the line. Predicted hinge residues are shown in grey. C) Single subunit of FH coloured according to eigenvector direction (positive as orange and negative as purple). Hinge residues are highlighted as grey. D) Molecular Dynamics simulations of hinge mutations shows altered hinge flexibility. Left: Schematic of the angle measured in each simulation, Right: Angle of WT (red), and R233H mutant FH (blue) over a 200 ns equilibrium molecular dynamics simulation.

We surmised that regions involved in the “hinging” of FH domains may influence the binding site assembly due to the proximity and reliance of the quaternary structure of multiple domains to make up the binding pocket. We used Gaussian Network Modelling (GNM) within prody(26, 27) to predict hinge residues between the subdomains of the protein (**Fig 2 B**). We used the second normal mode for calculation of hinge residues, as the first normal mode reflects an unphysiological bending of the centre of the protein. Calculating the hinge residues results in residues 196, 198, 232, 242, 270, 317, 401, 411, and 448 being the most likely “hinge points” in the structure, these residues are shown on a single subunit of FH, coloured by eigenvector direction in **Fig 2 C**. To assess how mutations to this region disrupt the quaternary structure of FH, we chose the simulate the known R233H mutant, and the wild type (WT) tetrameric assemblies for 200ns each using molecular dynamics simulations. Measuring the angles between CA atoms of two residues in the centre of the D2 and D3 regions with respect to the hinge reveals that the R233H mutant reduces the angle of the domains by an average of 8 degrees, and so leads to a partial occlusion of the catalytic site of FH (**Fig 2 D**). From this evidence we conclude that disruption of these hinges are likely to alter the binding site and assembly of FH – and are likely pathogenic. We chose to treat all mutations with CA atoms within 6 Å of any hinge residue as potentially LOF through disruption of the protein quaternary structure.

Overall, we infer that mutations near to either the binding site, or a hinge region of the protein are likely to disrupt or alter the protein function. We find that, from the FH mutation database, a significant proportion of mutations can be classed as either binding site-associated, or hinge-associated, including a number of known loss-of-function (LOF) variants. Whilst 42 residues in the 461 amino acid protein structure (9%) are classified as being “binding site-associated”, we find that 11 of the 30 (36%) known LOF mutations are within these residues, showing a clear bias towards binding site-associated mutations. Similarly, 55 of the 461 (12%) amino acids in the protein structure are classified as “hinge-associated”, and we find 7 of the 30 (23%) within the FH mutation database fulfil this classification, showing a lesser, but still large occurrence bias. Distance calculations for all potential mutations are included in **Table S1**.

### High-Throughput mutational stability screen of FH in silico

To study how mutations that are not near the binding site or hinge regions may have effects on the structure of the protein, we sought to generate predicted mutational energy changes (ΔΔG) for every potential amino acid substitution in the FH structure. We chose to use two conceptually different methods and use an average between the two methods to study each potential mutant. We used the FoldX method(10, 11), and the Rosetta cartesian_ddg method(12, 13), (hereafter described as the Rosetta method) to perform mutational energy calculations.

To perform mutant calculations, the pdb structure 5UPP was first relaxed using the FoldX RelaxPDB method, before each mutation and its resultant ΔΔG was calculated. We additionally calculate the Relative Solvent Accessible Surface Area (RSA) for each wild-type (WT) residue. Mutations on the surface of the protein are unlikely to dramatically alter the folding of the protein, so we chose to only consider a mutation potentially destabilizing if it is buried, defined as having an RSA <= 0.2.

We find a good agreement between the FoldX and Rosetta methods, with an r of 0.67 (p <0.0001) for all mutational energies (**Fig 3 A**). This correlation is good given previous reports showing an overlap between Foldx and Rosetta-ddg of only 12%-25% when considering stabilizing mutations (28). Notably however, both methods appear to agree on predictions of mutations with extremely high energy, but there is a significant portion of the distribution that shows a reasonably poor correlation, particularly mutations that have a predicted ΔΔG between 1 and −1 kcal/mol. We additionally chose a cutoff of 2.5 kcal/mol to classify mutations as destabilizing, as this provides a good separation between known loss of function and known benign mutations that are not binding site adjacent (**Fig 3 B**). Across all potential mutations we find that ∼45% (3968 out of 8778) meet this criterion (**Fig 3 C**). This fits roughly with historical data of mutational stability in T4 lysozyme, which found that 45% of mutational sites lead to structural inactivation of enzymatic function (29).

**Figure 3:**
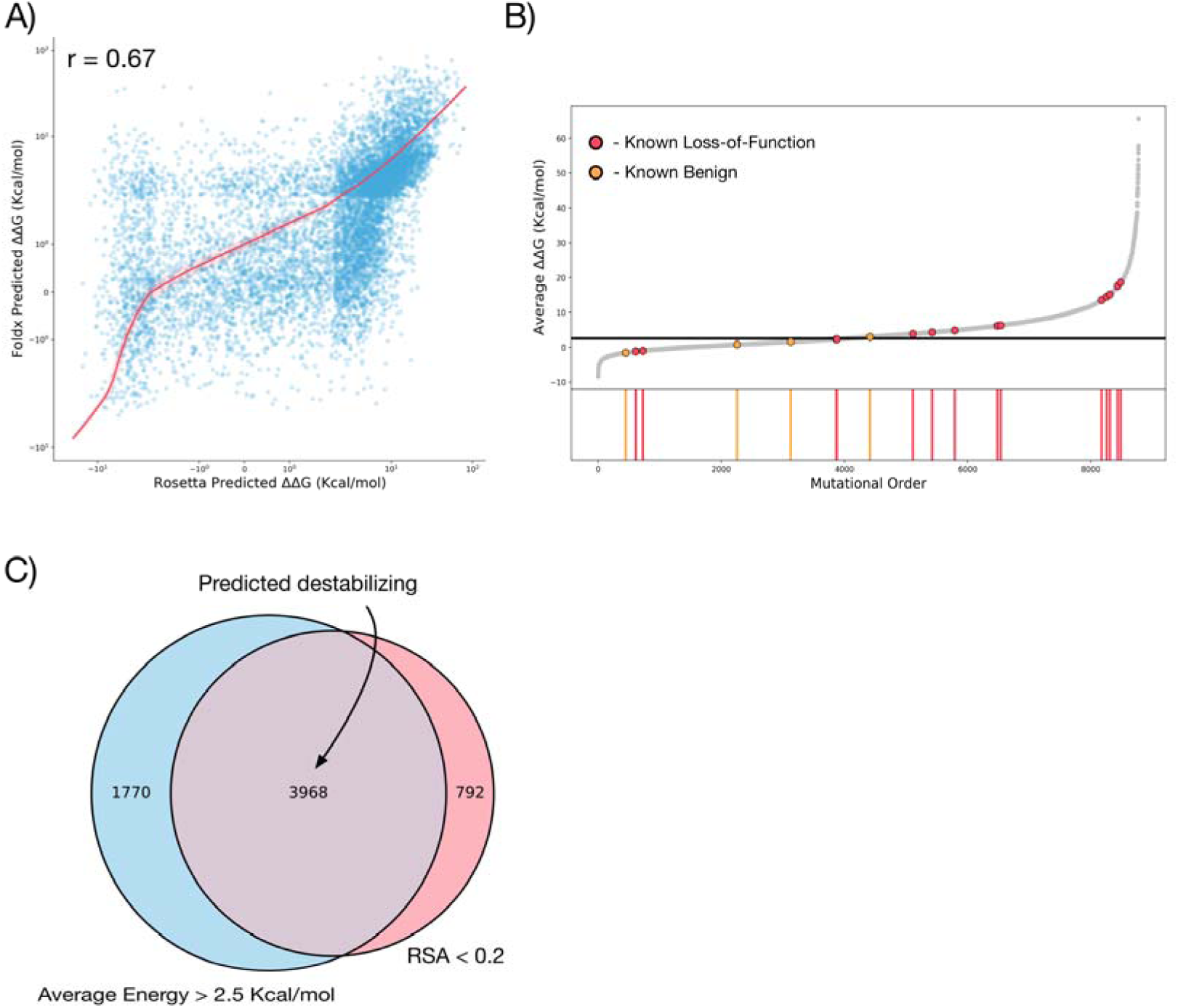
Prediction of Destabilizing Mutations. A) Comparison of ΔΔG calculations from FoldX and Rosetta. Correlation r is spearmans rank. B) Position of known loss-of-function (red) and benign mutations (orange) on the ΔΔG spectrum. Mutations are ordered in acsending mutational ΔΔG. Black line represents 2.5 Kcal/mol cutoff. C) Overlap between residues with a high predicted mutational energy (Defined as those with average ΔΔG > 2.5 Kcal/mol) and buried residues (RSA < 0.2). In total 3968 mutations are classified as destabilizing by taking the overlap between these two criterion.

Plotting mutational frequency for both methods, and their average for each residue (**Fig 4 A**) reveals that the most destabilizing mutations predicted by either method are in regions with a large number of buried amino acids, as expected. When plotting these mutations on the structure of the protein (**Fig 4 B**), we find the most significantly destabilizing mutations are those packed within the centre of D1, and on the interface between D1 and D2. This location suggests mutational disruption will alter the position of the D1/D2 interface, and thus will affect the binding site conformation, whereas mutations within the core D2 region are likely to influence the stability of the fully assembled tetramer.

**Figure 4:**
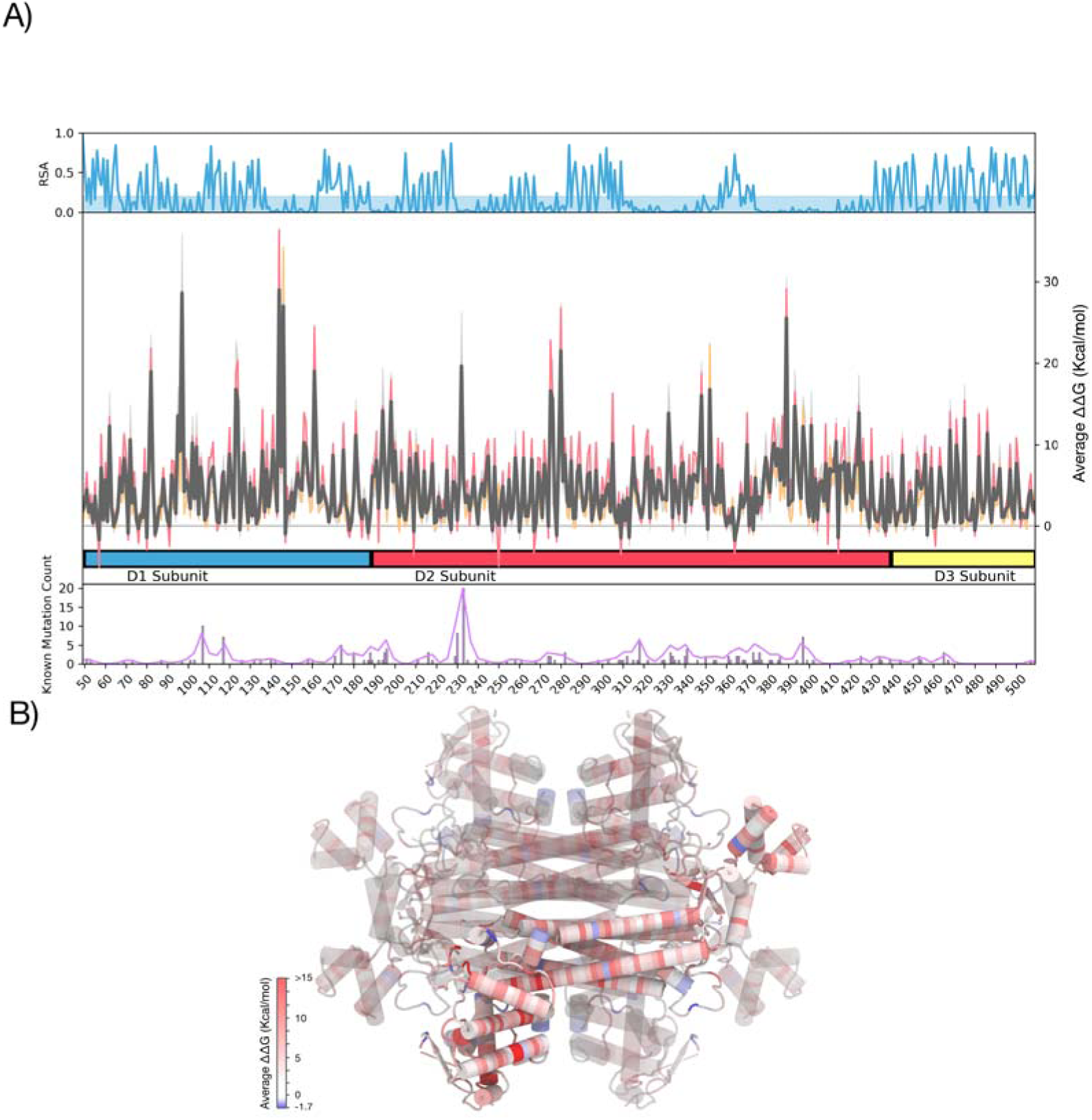
Predicted ΔΔG for every mutation in Fumarate Hydratase. A) Average mutational energy per residue in the FH structure. Top: Relative solvent accesible surface area (RSA) for every residue. Blue highlight indicates RSA < 0.2, classified as buried. Middle: average ΔΔG for each residue (Grey). Red and Orange lines represent average Rosetta and FoldX calculations respectively. Bottom: Mutational frequency from the FH mutation database for each residue in FH. B) Mutational ΔΔG applied to the structure 5upp of FH. Red indicates high average ΔΔG, and so represents areas where mutations are likely to disrupt the structure. Blue represents regions of generally stabilizing mutations.

### Existing mutations are accurately categorised based on known phenotypic effects

Overall, we define a scheme for classifying mutations into different categories of potentially disrupting, predicted LOF substitutions (**Fig 5 A**). The initial structure is relaxed using FoldX, before the binding site and hinge regions are calculated and classified, additionally mutations that are potentially destabilizing are defined based on average energy from the Rosetta and FoldX mutation methods, plus screened for buried mutations through calculating the RSA. This results in a categorisation for every mutation, where is it classified as predicted silent, binding site, hinge site, or destabilizing (including combinations of disruptive mutation types) (**Table S1**). Overall we classify 5811 out of 8778 (66%) mutations as potentially functionally disruptive, similar to a study of mutational effects on TP53, which found that roughly 50-60% of all possible mutations were functionally disruptive(30).

**Figure 5:**
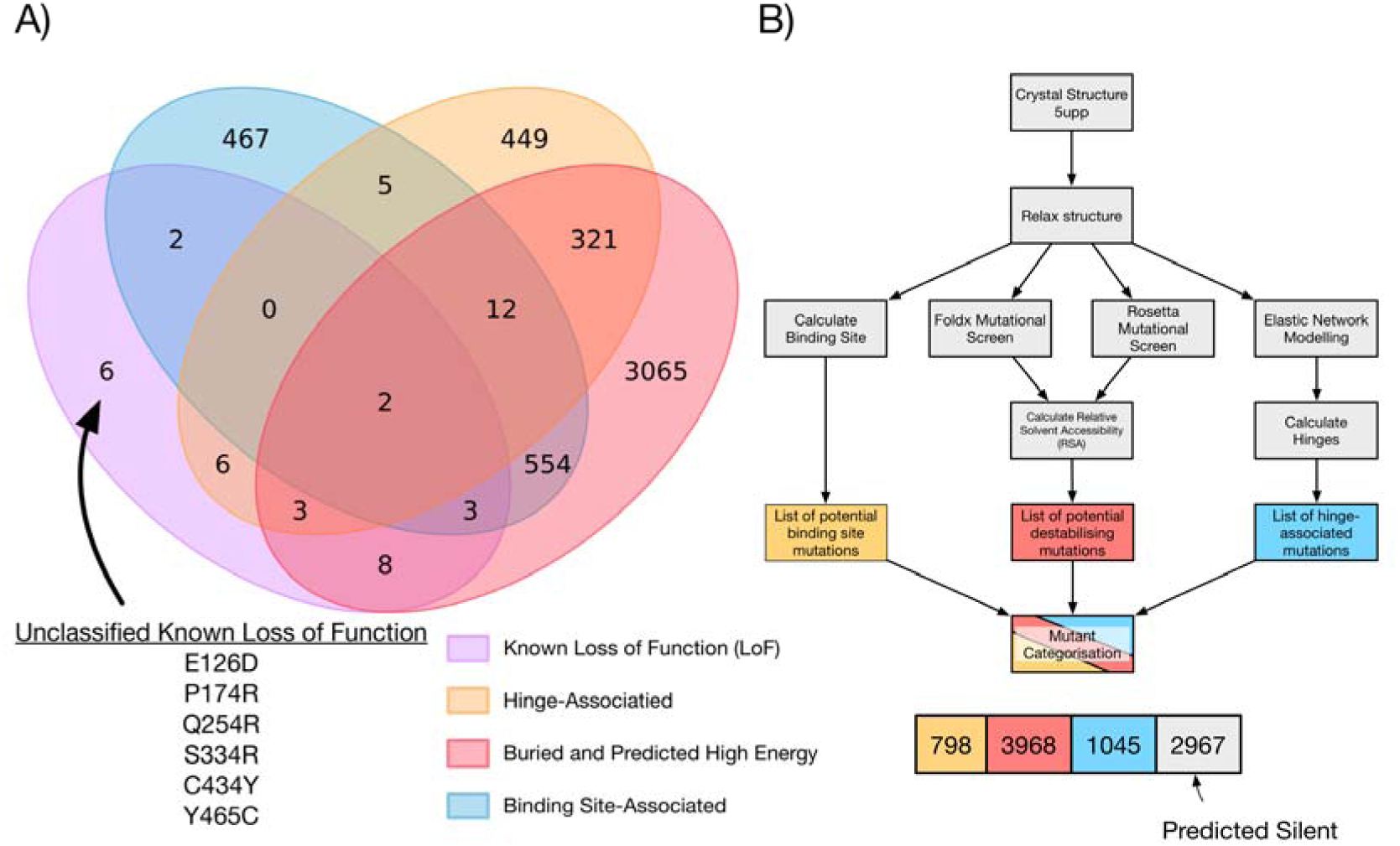
Prediction of known Loss-of-Function (LOF) mutations. A) Venn diagram showing overlap between Hinge-associated (orange), destabilizing (red) and Binding site-associated (blue) mutations. 30 known LOF mutations are included (purple) 24 mutations are correctly categorized as LOF, whilst 6 are incorrectly categorized as benign mutations. B) Schema for categorization of mutations in FH. The structure is initially relaxed using FoldX RelaxPDB, residues within 6Å of the binding site are calculated resulting in a list of 798 binding site associated mutations (orange). FoldX and Rosetta are used to calculate the ΔΔG for every mutation and this is subset by the relative solvent acessible surface area resulting in 3968 potentially destabilizing mutations (red). Elastic network modelling is performed to generate hinge regions of the protein, and residues within 6Å of hinges are calculated, resulting in 1045 hinge associated mutations (blue). 2967 mutations are predicted to be silent.

We compared our predictions with known mutations in the FH mutations database. We predicted a classification for all mutations within the database and compared this to a classification for each categorised as loss of function (LOF), benign, or of unknown functional effect. In total 34 mutations had a known (or implied) functional effect, whilst 73 were classified as unknown (**Table S2**). We find that 24 out of 30 (80%) mutations are correctly classified as LOF using our classification scheme, and 3 out of 4 (75%) are correctly classed as benign (**Fig 5 B**). Of the mutations incorrectly classified as benign when they are known to be LOF, two mutations involve cysteine (C434Y, Y465C), which is known to be modelled poorly by Rosetta cartesian_ddg, The mutation incorrectly classified as deleterious when it is listed as benign within the FH mutation database is R268G. We predict the R268G mutation to be both destabilizing (ΔΔG > 2.5 kcal/mol, RSA < 0.2) and hinge-associated. Whilst the mutation is listed as benign, no experimental information is cited, and PolyPhen-2(31), and Rhapsody also classifiy this particular mutation as damaging, indicating that the benign classification for this mutation may be questionable. We ran a molecular dynamics simulation of the R268G mutant. Simulations predict that mutant R268G reduces the hinge angle of the D1/D2 domains by ∼5 degrees (**Fig S1**), and supports previous evidence from the R233H mutant, that hinges within he protein can effect binding site assembly. Of the 73 unknown mutations, we predict that 28 are functionally benign, and 45 are potential LOF mutations.

### Mutations with unknown properties can be accurately predicted to be functional or neutral

To visualise all potential mutations in FH we chose to plot all mutations using umap(32). Umap attempts to cluster items by similarity, in a manner similar to prinical component analysis, or TSNE (33). We ran umap on all mutations using the 4 major axes involved in the classification – Minimum distance to a binding site residue, minimum distance to a hinge residue, average ΔΔG of mutation, and RSA for each residue (**Fig 6 A**). We find that distinct regions of the plot cluster into functionally different mutations when coloured by classification. There is a region specifically for hinge-associated mutations, binding site-associated, and unknown (not predicted damaging) mutations. In particular, the region of “unknown” (not classified as damaging) mutations overlaps significantly with a number of predicted destabilizing mutations, indicating that discrimination between these mutations is difficult, and perhaps not accurate with currently available data. We find that most of the benign mutations, aside from R268G are found clearly within the regions of predicted benign mutations. R268G clusters with the hinge mutation region as expected from our previous classification. For the known LOF mutations, we find they mostly cluster within the well defined regions for binding site, hinge, and destabilizing mutations. There are some mutations, particularly those which were misclassified, that fall within ambiguous regions of state space in the mutational landscape, and so are hard to classify using our defined criterion.

**Figure 6:**
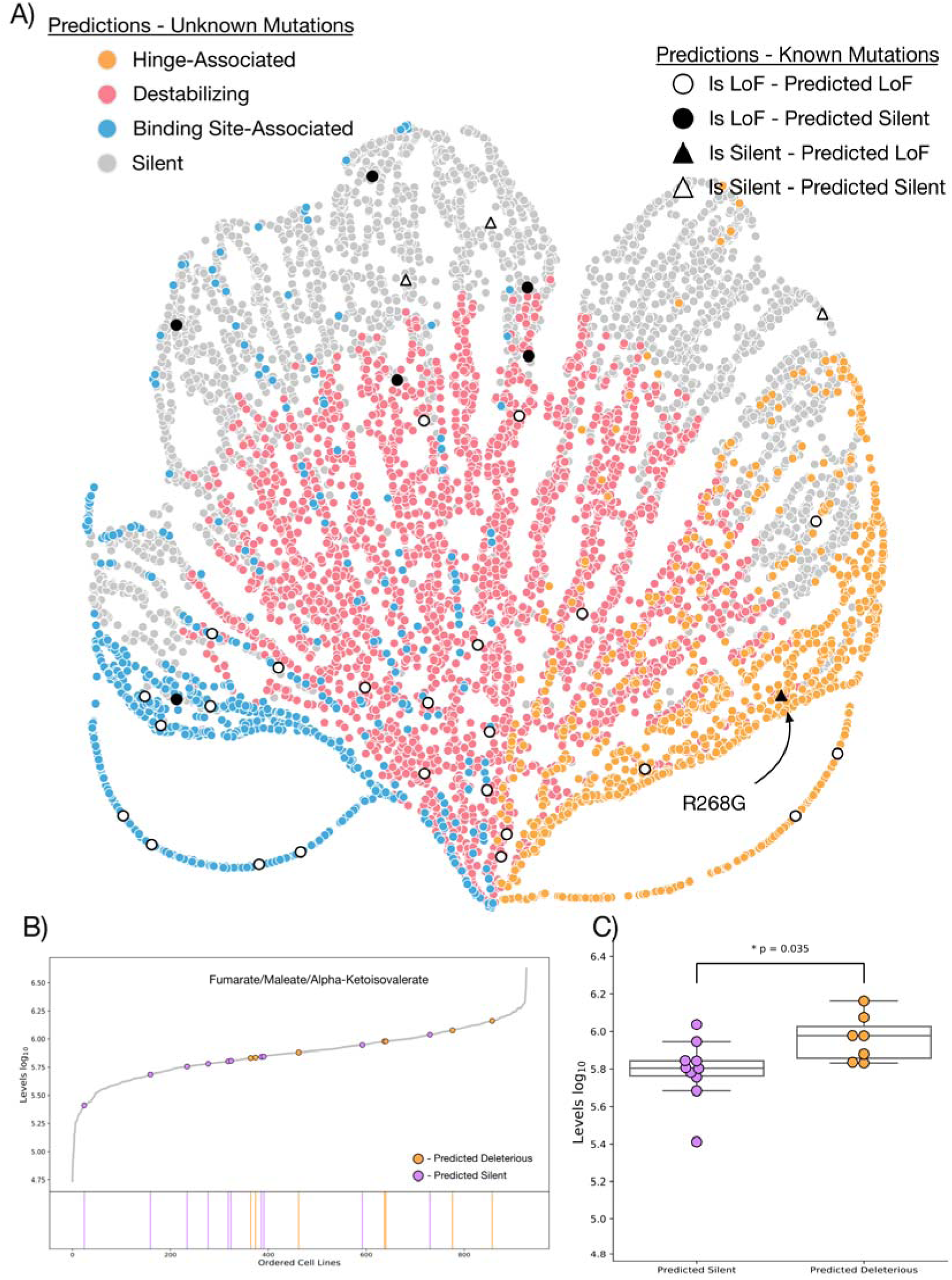
Mutational Landscape of Fumarate Hydratase. A) Umap for all mutations in FH. Mutations are coloured by classification. Hinge-associated (orange), Destabilizing (red), and Binding site-associated (blue) are shown clustered into groups. Predicted silent mutations (grey) are also shown. Overlayed are our predictions for characterized mutations in the FH mutation database. Mutations that are known Loss-of-Function (LOF) are circular and coloured according to whether we predict them to be LOF (black) or silent (white). Known benign mutations are in triangles, and also coloured according to whether we predict them to be LOF (black) or silent (white). The questionable known benign mutation R268G is labelled B) Mutations in the Cancer Cell Line Encyclopedia (CCLE) metabolomics data. All cell lines are ranked according to their detected levels of Fumarate/Maleate/Alpha-Ketoisovalerate. Coloured are cell lines with mutations in FH that we predict to bo LOF (orange), or silent (purple). C) Swarm plot for levels of Fumarate/Maleate/Alpha-Ketoisovalerate in mutant FH cell lines. Mutations predicted to be silent are significantly lower than mutations predicted to be LOF (p value represents independent T test).

To test the predictive power of our classification scheme we used the Cancer Cell Line Encyclopedia to look for changes in metabolite levels associated with mutations in FH(21, 34). We find 42 mutations (35 unique) in FH within 34 individual cell lines (**Table S3**). Selecting only for missense mutations yielded 25 mutations (20 unique) within 23 unique cell lines. We classified the mutations according to our criterion as either predicted LOF, or predicted benign. We find that by analysis of metabolomics data included in the CCLE database, mutations that we predict to be LOF have a higher average level of fumarate/mateate/alpha-ketoisovalerate detected in media than cells with predicted benign mutations (p = 0.035) – indicating that these cell lines may have an accumulation of fumarate as a result of inactive levels of FH (**Fig 6 B,C**).

## DISCUSSION

In conclusion, we have shown, using a comprehensive combination of techniques, that we can categorise accurately the functional effects of any potential missense mutation in FH. Beyond FH, we present an integrated series of methods that can be adapted for mutationally screening any protein for functionally relevant mutations in a reasonably small amount of computational time. Our workflow predicts the functional effects of all mutations that can be compared to existing methods based on machine-learning principles such as rhapsody and polyphen, at significantly lower time and effort expenditure than experimental characterization. Whilst some other methods incorporate some manner of structural analysis in their predictions, ours demonstrates a new perspective, as it explicitly models every potential mutation in a structure, allowing it to interface directly with other computational techniques in the field such as molecular dynamics simulations to further study mutations of interest.

Biologically we propose three ways in which mutations can potentially disrupt the catalytic activity of FH, in particular we find that addition of hinge altering mutations are necessary for classification of many known LoF mutations, indicating that there is a biological relevance, and hinting at a mechanism for, mutations that change the flexibility and stiffness of protein hinges in this case. Additionally, we chose to exclude site B from our analysis of mutation disruption and find that we are able to classify almost all known mutations without its inclusion. This implies that mutations in site B may not have functional or disease-related relevance, despite some evidence that site B can alter catalytic activity of the enzyme(35). This is reinforced by the fact that 27 of the 461 residues within the protein structure are classified as near site B (6%), and only 3 of 30 residues in the FH mutation database (10%) are near to site B, showing a poor to negligible enrichment of mutations in site B when compared to similar calculations for site A.

Fumarate hydratase represents a good first-use case for high-throughput mutational screen due to the need to understand mutations in their functional context, but as mutational detection techniques, and high-throughput mutational studies increase the need to be able to classify mutations confidently as benign and LOF is more important. Here we show that our method accurately classifies known LOF and benign mutations with a high degree of accuracy, and predict which mutations discovered in the human population are likely to have functional relevance, and therefore predispose patients to particular metabolic diseases.

Whilst the accuracy of our method with the current data is high, there are clear regions where the analysis is not able to discriminate between mutations on the borderline between destabilizing and benign, this results from the lack of accuracy in the mutational ΔΔG calculations, despite using the best available methods at time of study(13). As better methods become available it will be of interest to improve upon this work to attempt a more accurate classification.

Finally, whilst the work here focusses on a single molecule within the TCA cycle, FH, structural data has existed for a large number of enzymes within the cycle for some time(36–39), and it would be of great interest to look into mutations across entire metabolic pathways. With this study laying the groundwork, it will be of future interest to model all mutations in all enzymes, and attempt to further link these with genomic and metabolomic data that is already available. Whilst the computational intensity of this work is high, it is feasible to screen thousands of mutations in only a few days of computational time.

## METHODS

All data used in this study, including the code used in generating all figures from raw data is available publicly at: https://github.com/shorthouse-mrc/Fumarate_Hydratase

### FH mutation database

The FH mutation database was downloaded from the Leiden Open Variation Database(9) (https://databases.lovd.nl/shared/variants/FH/unique). Missense mutations were manually curated into categories (Loss of Function, Benign, and Unknown) based on their implied clinical classification, and variant remarks, which contained information regarding FH enzymatic activity.

### Mutational Clustering

Mutational clustering was performed with the NMC clustering algorithm, which attempts to discern the likelihood of a mutation spectrum occurring by random chance. NMC returns clusters of mutations that are statistically significant. We chose to run the NMC algorithm using the R library iPAC(25), using an alpha cutoff value of 0.05, and the Bonferroni multiple test correction method (see supplementary code).

### Gaussian Network Modelling

GNM was implemented using the Prody package in python(40). A Kirchhoff matrix was constructed using the gnm.buildKirchhoff command with the parameters cutoff = 10.0 and gamma = 1.0. Normal modes were then calculated using the gnm.calcModes() command. Predicted hinges were assessed using the gnm.getHinges() command. The full protocol, implemented in a jupyter notebook is available at https://github.com/shorthouse-mrc/Fumarate_Hydratase.

### Molecular Dynamics Simulations

Molecular dynamics was performed using Gromacs version 2018.1(41). We chose to simulate proteins using the GROMOS 54a7 forcefield(42).

The protein structure was first repaired using FoldX(10) “RepairPDB” with the following command:

$foldx --command=RepairPDB --pdb=5upp.pdb --ionStrength=0.05 --pH=7 -- vdwDesign=2

The protein was then placed in a cubic box size 15 x 15 x 15 nm and solvated with roughly 90,000 spc water molecules. Counterions were introduced to a neutral charge, and to a concentration of 0.05 mol/litre. The system was energy minimized using the steepest descents algorithm until the maximum force,F_max_, of the system reached below 1000 kJ/mol/nm.

Equilibration was performed using the NVT, followed by the NPT ensembles for 100 ps each. We chose to use the verlet cutoff scheme and PME electrostatics, and utilized periodic boundary conditions in the x,y, and z planes.

Molecular dynamics was performed for 200 ns retaining velocities from the NPT equilibration. We used the V-rescale temperature coupling scheme, and Parrinello-Rahman isotropic pressure coupling.

### FoldX ΔΔG Calculations

FoldX predicted ΔΔG was calculated using the PositionScan command within FoldX4. Positionscan was run on each residue in the protein structure sequentially using the following command:

$foldx --command=PositionScan --pdb=5upp.pdb --ionStrength=0.05 --pH=7 -- vdwDesign=2 --pdbHydrogens=false --positions=49

For positionscan on the 49^th^ residue (I think residue in the FH structure 5UPP).

### Rosetta ΔΔG Calculations

Rosetta predicted ΔΔG was calculated using the cartesian_ddg method as described in Kellogg et al:

$path/to/source/bin/cartesian_ddg.static.linuxgccrelease -in:file:s 5upp.pdb - in::file::fullatom -database /path/to/database/ -ignore_unrecognized_res true - ignore_zero_occupancy false -fa_max_dis 9.0 -ddgccartesian -ddg::mut_file mutfile.txt -ddg::iterations 3 -ddg::dump_pdbs true -ddg::suppress_checkpointing true -ddg::mean true -ddg::min true -ddg:output_silent true -bbnbr 1 - beta_nov16_cart > logfile.log

ΔΔG was calculated by averaging the energy of 3 models of each mutation and comparing it to the WT calculation.

### Umap

We used Umap(32) based on the github repository at www.github.com/lmcinnes/unmap

### Cancer Cell Line Encyclopedia Data

Cancer Cell Line Encylopedia (CCLE) mutation data was downloaded from the Broad Institute at: https://portals.broadinstitute.org/ccle/data. Metabolomics data was obtained from the supplementary data of Li et al(21).

### Data Analysis

Both MDanalysis(43) and Biopython(44) were used for analysis of structural data. Data analysis workflows are available in jupyter notebooks available at https://github.com/shorthouse-mrc/Fumarate_Hydratase.

## AUTHOR CONTRIBUTIONS

DS and BAH conceived the study and wrote the manuscript. DS generated all data and performed all analysis. All authors were responsible for editing of the manuscript.

## ACKNOWLEDGEMENTS

We thank the Frezza group, in particular Christian Frezza for support and constructive feedback during the generation of this manuscript. This work was supported by the Medical Research Council (grant no. MR/S000216/1). M.W.J.H. acknowledges support from the Harrison Watson Fund at Clare College, Cambridge. B.A.H. acknowledges support from the Royal Society (grant no. UF130039).

