## Supplementary Text for "The Landscape of Mutations in Human Fumarate Hydratase"

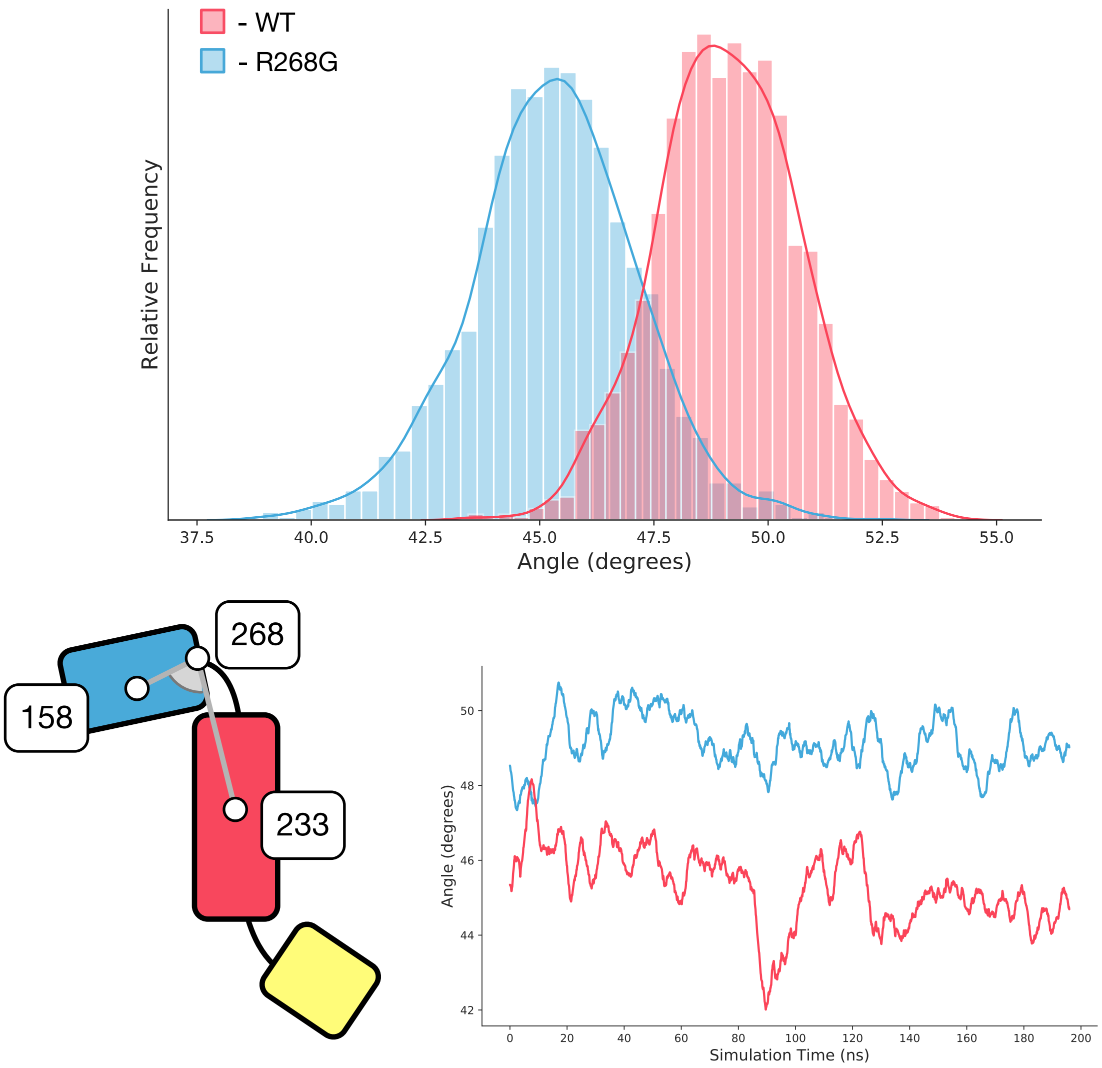


Figure S1: Molecular Dynamics of WT (red) vs R268G Mutant forms of FH (blue). Shown is the angle between residues 158, 268, 331.

Table S1: Classification for all potential mutations in FH
Table S2: Prediction for all mutations in the FH database
Table S3: Prediction for all mutations in FH in the Cancer Cell Line Enyclopedia
